# Human visual cortex is organized along two genetically opposed hierarchical gradients with unique developmental and evolutionary origins

**DOI:** 10.1101/495143

**Authors:** Jesse Gomez, Zonglei Zhen, Kevin S. Weiner

## Abstract

Human visual cortex is organized with striking consistency across individuals. While recent findings demonstrate an unexpected coupling between functional and cytoarchitectonic regions relative to the folding of human visual cortex, a unifying principle linking these anatomical and functional features of cortex remains elusive. To fill this gap in knowledge, we combined independent and ground truth measurements of human cytoarchitectonic regions and genetic tissue characterization within the visual processing hierarchy. Using a data-driven approach, we examined if differential gene expression among cortical areas could explain the organization of the visual processing hierarchy into early, middle, and late processing stages. This approach revealed that the visual processing hierarchy is explained by two opposing gene expression gradients: one that contains a series of genes with expression magnitudes that ascend from the first processing stage (e.g. area hOc1, or V1) to the last processing stage (e.g. area FG4) and another that contains a separate series of genes that show a descending gradient. In the living human brain, each of these gradients correlates strongly with anatomical variations along the visual hierarchy such as the thickness or myelination of cortex. We further reveal that these genetic gradients emerge along unique trajectories in human development: the ascending gradient is present at 10-12 gestational weeks, while the descendent gradient emerges later (19-24 gestational weeks). Interestingly, it is not until early childhood (before 5 years of age) that the two expression gradients achieve their adult-like mean expression values. Finally, additional analyses in non-human primates (NHP) reveal the surprising finding that only the ascending, but not the descending, expression gradient is evolutionarily conserved. These findings create one of the first models bridging macroscopic features of human cytoarchitectonic areas in visual cortex with microscopic features of cellular organization and genetic expression, revealing that the hierarchy of human visual cortex, its cortical folding, and the cytoarchitecture underlying its computations, can be described by a sparse subset (~200) of genes, roughly one-third of which are not shared with NHP. These findings help pinpoint the genes contributing to both healthy cortical development and the cortical biology distinguishing humans from other primates, establishing essential groundwork for understanding future work linking genetic mutations with the function and development of the human brain.

## Introduction

One of the most reproducible findings across species in biology and neuroscience is the identification and definition of the visual processing hierarchy. Indeed, despite debates regarding the exact number and definition of areas, as well as the serial or non-serial nature of this hierarchy, researchers acknowledge the existence of early (e.g. striate cortex, or V1), middle (e.g. areas in extrastriate cortex), and late (e.g. areas in inferior, or ventral, temporal and lateral occipitotemporal cortices) processing stages of the visual hierarchy^1–3^. In humans, the visual processing hierarchy is identifiable with a multitude of methods in both living brains and postmortem tissue. For example, areas composing the visual processing hierarchy have been identified in living brains using anatomical and functional magnetic resonance imaging^2–6^, as well as in postmortem brains based on cytoarchitecture^7–13^, myeloarchitecture^14,15^, and receptor architecture^16–18^. Interestingly, and contrary to classic findings^19–22^, recent research in human postmortem brains indicates a tight correspondence between cellular transitions of cytoarchitectonic areas and cortical folding across the visual hierarchy^13,23–25^. Similarly, recent research also indicates a tight correspondence between functional regions and cortical folding across the visual hierarchy for early^26^, middle^27–29^, and late^23,24,30^ visual regions. In terms of the latter, this striking consistency is conserved across development^30–34^, and is causally implicated in different aspects of visual perception^35–39^. Given this tight relationship among cellular organization, functional organization, cortical folding, and perception, the visual processing hierarchy of the human brain is an ideal test bed to ask a fundamental, yet unanswered, question in neuroscience: *What is the shared principle by which brain function and structure are linked that results in the shared brain organization and behaviors across individuals?*

When faced with this question, one might quickly intuit that a feasible underlying answer is likely genetic in nature. Indeed, given recent results showing that genetics play a role in the broad hierarchical division of cortex - for example, differentiating primary sensory areas such as motor cortex from association areas in prefrontal cortex^40^ - it seems likely that genetics may also play a role in determining the fine-scale hierarchical division of human visual cortex. While measuring the expression of genes across discrete regions of cortex *in vivo* is infeasible, recent advances in transcriptomics and brain mapping have resulted in public databases detailing the transcriptome across the human cortical surface^41,42^. Furthermore, previous work from our group has provided multimodal solutions for linking different types of data to one another across spatial scales - for instance, linking functional regions at a millimeter scale in individual living human brains to cytoarchitectonic organization at a micron scale in individual postmortem brains^25,30,43^. Thus, by applying this multimodal approach to gene transcription and cytoarchitectonic data, we have the unprecedented opportunity to ask what role genes play in the areal differentiation of the human visual processing hierarchy based on cytoarchitecture.

To fill this gap in knowledge, we employed the Allen Human Brain Atlas, which is a transcriptomic analysis of cortical tissue sampled throughout the neocortex in six brains, resulting in cortical surface-based maps of the expression magnitude of over 20,000 genes. Expression maps were aligned to a common anatomical space where they could be averaged across subjects and hemispheres (see Materials and Methods). To examine what role genes play in differentiating stages of the visual processing hierarchy, we aligned 13 cytoarchitectonic regions of interest (cROIs) from 20 postmortem hemispheres (maximum probability maps from the Jülich atlas^7–9,11–13^: https://jubrain.fz-juelich.de) to the same anatomical surface. By isolating those genes with the most significant differential expression across the 13 cROIs, we found that the visual processing hierarchy is described by two opposed genetic expression gradients: an ascending gradient that increases its expression along the hierarchy, and a descending gradient that decreases its expression magnitude along the hierarchy. We replicate these findings in a separate dataset, as well as provide developmental and evolutionary insight into these gradients by leveraging additional transcriptome measurements from developmental (e.g. gestational to 60 years of age) and macaque datasets. Repeating our analyses with these additional data reveals that the ascending and descending genetic gradients emerge at different points of human development (10-12 versus 19-24 gestational weeks) and only the ascending gradient is present in macaques. These findings (a) empirically support that the consistent organization of the human visual processing hierarchy is achieved through opposed genetic expression gradients, as well as (b) begin to establish a biological model that links measurements at the levels of genes and cellular organization with the macroscopic features of the human visual processing hierarchy.

## Results

### Opposed genetic gradients contribute to cytoarchitectonic divisions of human visual cortex

To identify those genes that contribute to areal differentiation of the visual processing hierarchy in human ventral and lateral occipitotemporal cortices, we assessed gene expression profiles that were significantly different across cytoarchitectonic regions of interest (cROIs). In brief, cROIs were defined previously^7–13^ in individual postmortem brains using the cell-density profile across layers of cortex (Fig 1A). Transition points from one cytoarchitectonic region to another were defined by an observer-independent algorithm that was blind to cortical folding. Once cROIs were defined in individual participants, they were projected from 2D histological slices to 3D cortical surface reconstructions, allowing all cROIs from each individual to be aligned using cortex-based alignment to a common space (e.g. the MNI152 average brain). Maximum probability ROIs describing the most consistent location of a given cROI across postmortem brains are shown in Figure 1B. Within each cROI, there were multiple tissue samples (sample numbers by cROI shown in Fig 1C), each of which was submitted to DNA microarray analysis to quantify the expression of 20,737 genes. For each gene, we tested (e.g. with an ANOVA as done in recent work also linking cytoarchitectonic and transcriptomic data, but in frontal cortex^44^) if the expression levels from tissue samples were significantly different across cROIs. To equalize tissue sample numbers per ROI for the ANOVA, we grouped the cROIs into four groups (Fig 1D). A histogram (Fig 1E) of resultant *p* values (negative log-transformed) for each gene revealed that over 50% of quantified genes are not significantly different in their expression levels across cROIs (all *p*s > 0.05). In order to target those genes that most strongly contribute to areal differentiation and to correct for multiple comparisons, we restricted our further analyses to only the top 1% (n=200) of genes that were most differentially expressed.

**Figure 1:**
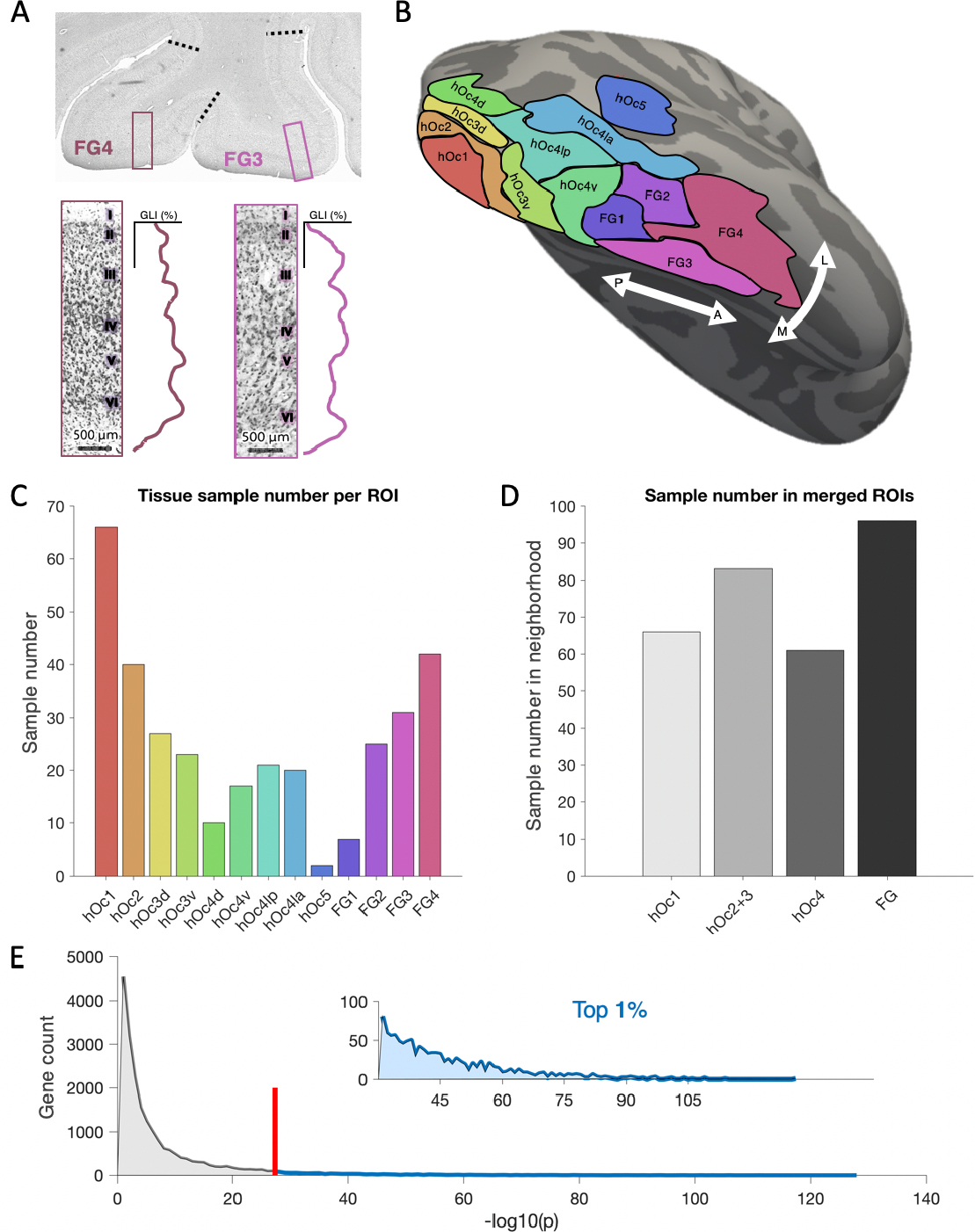
Tissue samples within each cytoarchitectonic area and merged cortical neighborhoods. **(A)** A schematized version of how cytoarchitectonic areas were defined using an observer-independent algorithm in previously published work (please see Amunts & Zilles 2001 for quantification details). Example windows taken from histological stains of human visual cortex, detailing the gray-level index (GLI) along the cortical depth. Gray-level indices (GLI) were measured within these windows which slid along the cortex, and boundaries (black-dotted line) were drawn when GLI profiles changed significantly along several dimensions. The image of the histological section provided by Evgeniya Kirilina. GLI profiles adapted from Weiner et al., 2017. **(B)** The cytoarchitectonic regions of interest (cROIs) delineated on the FreeSurfer cortical average surface. We considered 13 cROIs: human Occipital 1 (hOc1), hOc2, hOc3 dorsal (hOc3d), hOc3 ventral (hOc3v), hOc4 dorsal (hOc4d), hOc4 ventral (hOc4v), hOc4 lateral posterior (hOc41p), hOc4 lateral anterior (hOc41a), hOc5, Fusiform Gyrus 1 (FG1), FG2, FG3, and FG4. **(C)** Once the cROIs were aligned to the transcriptomic data (Materials and Methods), we quantified the number of tissue samples from the 6 postmortem brains included in the Allen Human Brain Atlas (AHBA) that fall within each cROI. The colors for each cROI are the same as in B. **(D)** Prior to gene selection, we first grouped gene expression samples into four cortical neighborhoods (differently shaded gray bars) as there was an unequal number of tissue samples in each cROI. **(E)** In order to select genes for further analyses, we ran a 1-way analysis of variance (ANOVA) with cROI group as a factor. Histogram of the resulting p-values (negative log transformed) from the ANOVA. We ranked the genes according to the p-values from the ANOVA and selected the top ~1% (200 genes; beyond the red line and shown in blue) of genes for further analysis. This thresholding approach was taken in order to avoid including genes that may result in significance simply as a function of multiple comparison. Inset illustrates a zoomed-in histogram of the top 1% of genes.

To gain a better understanding of the gene expression patterns across cytoarchitectonic regions, the top 200 genes were submitted to an agglomerative hierarchical clustering algorithm (see Materials and Methods for details). Surprisingly, genes clustered predominately into two groups at the highest level, whose branch distance dwarfed all other clustering levels (Fig 2A). As a control analysis, we repeated this clustering algorithm on the bottom 200 genes (smallest 200 *p* values from the gene histogram). This analysis revealed that this binary split was unique to the top 200 genes, as the bottom revealed a much more homogenous branching pattern across cluster levels (Fig 2A). We next produced a raster plot of gene expression magnitude (z-score normalized) to examine the expression patterns driving the split between the top genes. Strikingly, the two gene clusters are organized into two groups defined by opposed expression gradients along the visual processing hierarchy. In Figure 2B, the areas are ordered based on the number and anatomical location of the area; for example, hOc1 is first, then hOc2, and so forth, while FG4 is last and comes after FG3. One group forms a descending gradient whose composite genes express most highly in hOc1 (e.g. striate cortex) and decrease as one ascends the hierarchy across extrastriate cortex to the last area, FG4. The other group forms an ascending gradient with increasing expression levels up the hierarchy from hOc1 to FG4. Proportionally, there is a greater number of genes contributing to the ascending gradient (two thirds) compared to the descending gradient (one third). For a list of the ascending and descending genes, we ask the reader to refer to Supplementary Table 1. We additionally confirmed the existence of these gradients using a data-driven approach by submitting all gene expression profiles (not just the top 1%) of the 13 cROIs to a principal component analysis, which verified that the first component captures ascending and descending expression patterns within the most differentially expressed genes (Fig. S1).

**Figure 2:**
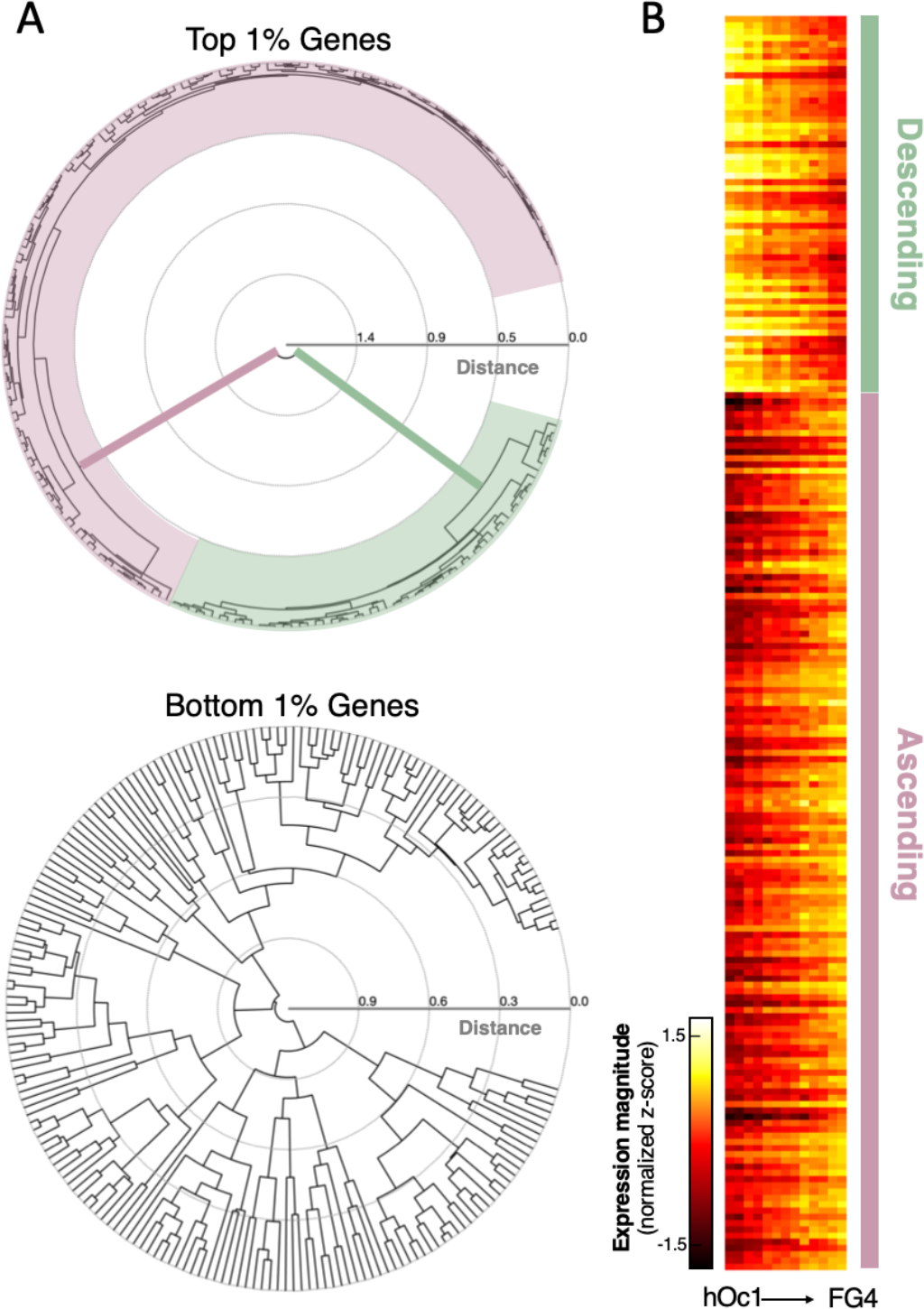
Unique binary clustering of the top 1% of genes reveals opposed expression gradients in human visual cortex. **(A)** Dendrograms showing the algorithmic clustering of the top and bottom 1% of genes. Top 1% of genes cluster predominantly into two groups at the highest level, colored here in pink and green. **(B)** A 200 × 13 matrix in which each column is an ROI arranged according to position within the visual processing hierarchy (e.g. hOc1 on the left) and each row is a gene. The first level of clustering of the top 200 significant genes showing differential expression reveals two distinct groups of cytoarchitectonic ROIs in human visual cortex: 1) a group with a descending gradient (green) in which genes have the highest expression in early visual cortex (e.g., hOc1, hOc2, etc.) and decrease their expression across the visual processing hierarchy and 2) an ascending gradient (pink) in which genes have lower expressions in early visual cortex and show increasing expression levels into high-level visual cortex. Expression levels are normalized to the maximum expression level (reads per kilobase million) across all tissue samples, and warmer colors indicate higher expression levels, while darker colors indicate lower expression levels.

### Clustering the expression levels of the ascending and descending gradients produces an ordering of cytoarchitectonic regions reflective of the human visual processing hierarchy

Are the expression patterns of this sparse subset of genes capable of ordering the hierarchy of cytoarchitectonic regions? That is, we ask if the ascending and descending gradient gene expression levels can discriminate distinct clusters of cROIs from early areas of the hierarchy (e.g. hOc1, hOc2, and hOc3) to middle areas of the hierarchy (e.g. the hOc4 areas and hOc5), and finally, to late areas of the hierarchy (e.g. FG1 through FG4). To answer this question, we produced average expression patterns within the descending and ascending gradient gene clusters. As expected, clusters formed either a linearly decreasing (Fig 3A) or increasing (Fig 3B) gradient across the visual processing hierarchy from hOc1 (earliest stage) to FG4 (latest stage). We submitted the expression patterns of the top 200 genes (averaged across all tissue samples within a cROI) from the thirteen cROIs to an agglomerative clustering algorithm. Before clustering, cROIs were ordered according to the Euclidean distance of their expression profile from that of hOc1, such that the ordering of the x-axis in the resulting dendrogram meaningfully reflects inter-regional distances from hOc1. As such, this rooted-leaf dendrogram has clusters that cannot be rotated at the lowest level as in other dendrograms. At its highest level, the resulting dendrogram clusters early (hOc1, hOc2, hOc3v, and hOc3d) from mid and late visual areas (hOc4v, hOc4d, hOc41p, hOc41a, hOc5, FG1, FG2, FG3, and FG4). At the next cluster level, hOc1 and hOc2 occupy a cluster separate from that of hOc3d and hOc3v. The next closest cluster contains the hOc4 regions, followed by hOc5 and FG1 and FG2. The last cluster, and the most distant from hOc1, contains FG3 and FG4. We emphasize that the clustering within this dendogram is not just reflective of macroanatomical proximity. For example, hOc3d and hOc3v are within the same subcluster and yet, are located centimeters apart in cortex. Likewise, hOc4v is located within the posterior transverse collateral sulcus in ventral occipitotemporal cortex^25^, while hOc4d is located within the transverse occipital sulcus on the lateral surface of the brain^10^, and yet, both areas are positioned in the same subcluster. Thus, macroanatomical proximity is not driving the ordering of areas within the dendogram. Finally, to evaluate the probability of reproducing the exact ordering of the visual hierarchy by chance, we used a bootstrap approach (n=10,000; see Materials and Methods), which shuffles the gene expression profile within each cROI on each bootstrap. When implementing this bootstrapping approach, we find that the ability of the top genes to correctly order the cROIs is highly significant (p<0.00001).

**Figure 3:**
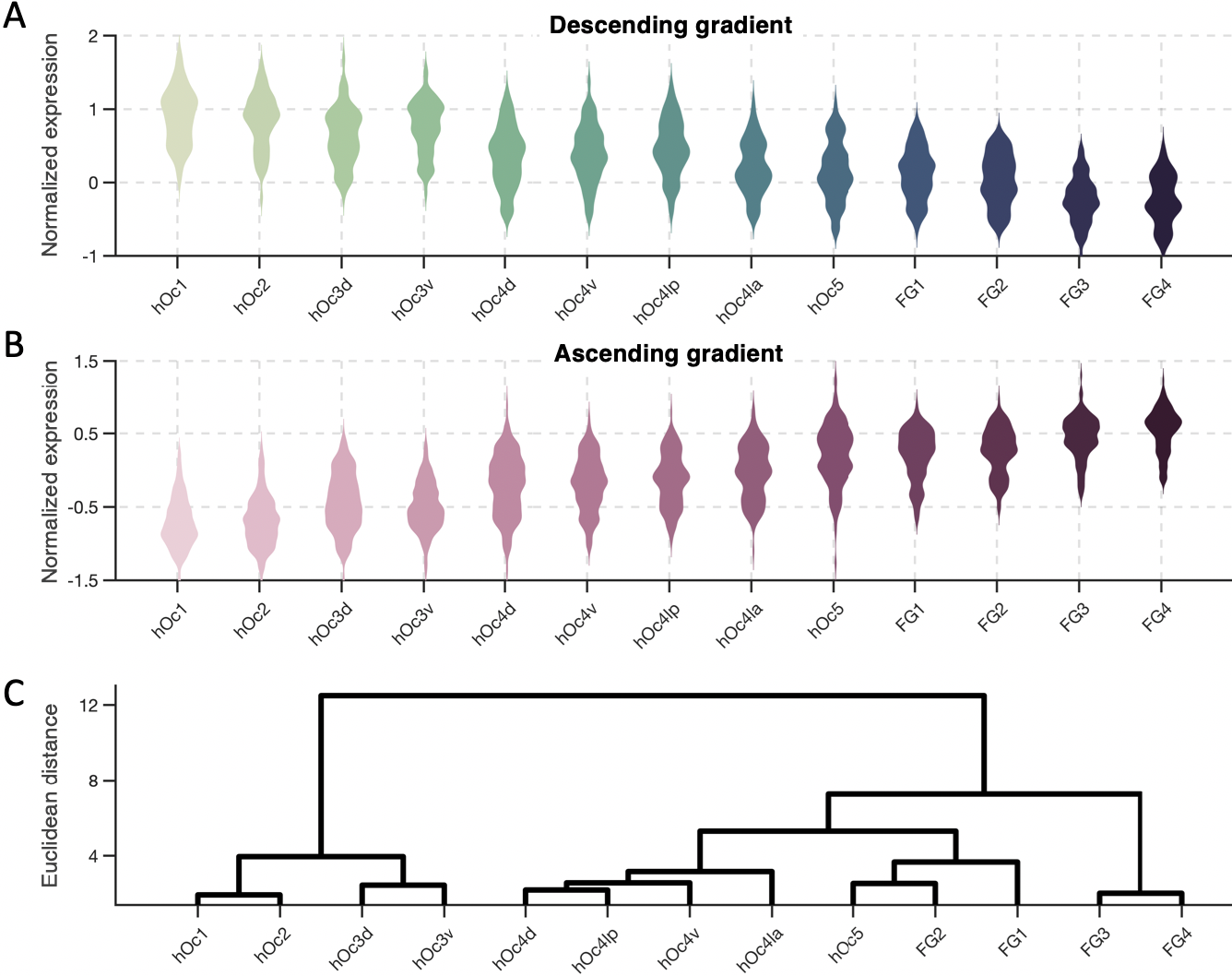
Average expression levels of the ascending and descending genetic gradients reproduce the organization of the visual processing hierarchy. (A) Gene expression levels (z-score normalized) shown across cytoarchitectonic regions for descending gradient genes. Color shade denotes relative hierarchy level (higher levels are shown in darker colors). (B) Same as A, but for ascending gradient gene expressions. (C) Dendrogram produced from a clustering algorithm on expression levels from the two gene clusters. The ordering of cROIs along the x-axis in the dendrogram is meaningful, as it represents the distance from hOc1 at the level of dendrogram leaves. Consequently, this dendrogram has rooted leaves and the clusters at this lowest level are not rotatable as in other dendrograms (Materials and Methods). This data-driven approach successfully reproduces the visual processing hierarchy by using the genetic expression levels of the two opposing gradients.

### Opposed genetic gradients correlate with cortical thickness and myelination

To help elucidate the cortical mechanisms that are influenced by these opposed gene gradients, we next asked what anatomical features of the cortex covary with expression patterns across the visual hierarchy. We specifically focus on two anatomical measures: myelination and cortical thickness. We focus on the former given previous findings demonstrating that V1 (hOc1) is a primary sensory region that is heavily myelinated at birth^45^, while higher-level visual cortex demonstrates delayed and protracted myelination^30^. We focus on the latter given previous findings demonstrating that the thickness of the cortical sheet^46^ distinguishes early sensory regions from regions located within higher-level association cortex. We used data from the Human Connectome Project (HCP)^47^, which is a large database containing MRI scans from 1096 adults and provides T1-weighted (T1w) and T2-weighted (T2w) scans that can be used to derive both measures of cortical thickness and a proxy of cortical myelin content (e.g. the ratio of T1w/T2w scans, which is a measure of tissue contrast enhancement related to myelin concentration, see Materials and Methods). We thus aligned individual subject maps of cortical thickness and maps of the ratio of T1w/T2w using cortex-based alignment to the FreeSurfer cortical average to which the cROIs were also previously aligned using a similar procedure^25,30^. Then, for each cROI, we computed the average thickness or T1w/T2w ratio across all HCP subjects. Illustrated in Figure 4A, we find that the T1w/T2w ratio produces a descending gradient as one ascends the visual processing hierarchy. Cortical thickness, in contrast, produces an ascending gradient along the visual hierarchy. An ANOVA with grouping variables of gradient type (T1w/T2w, thickness) and cROI reveals significant main effects (F’s_(12,28470)_>50, P’s<0.000001) and a significant interaction (F_(12,28470)_>50, P<0.000001). Cortical flat maps of average T1w/T2w ratio and cortical thickness from HCP subjects are illustrated in Figure 4B.

**Figure 4:**
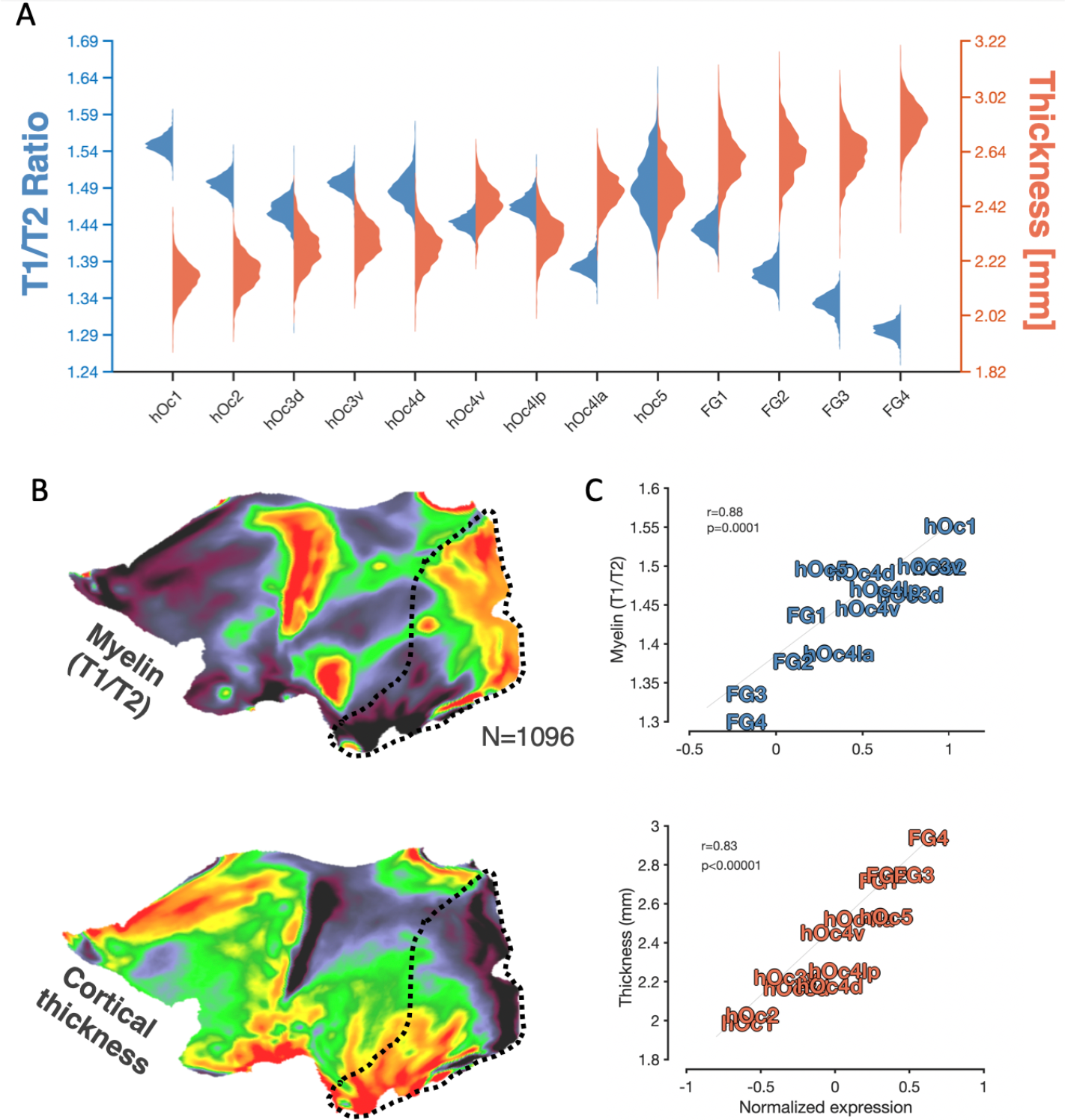
Positive- and negative-gradient genes mirror inter-areal differences in cortical myelination and thickness. (A) Violin plots depicting the distribution of tissue contrast enhancement (e.g. a ratio of T1w/T2w scans, which is a metric reflective of tissue contrast enhancement associated with myelin content) in blue, and cortical thickness in orange, across HCP subjects (n=1096) in cytoarchitectonic ROIs of the visual processing hierarchy. (B) Cortical flat maps depicting the average T1w/T2w ratio or cortical thickness across HCP subjects. Black-dotted outline highlights visual cortex. (C) Top: Correlation plot between the average “myelin” content (T1/T2 ratio) in cROIs and average expression magnitude of genes included within the descending gradient. Bottom: Same as above but for the average cortical thickness of a given cROI and and expression magnitude of genes included within the ascending gradient.

To see if cortical myelination is significantly correlated with the expression levels of genes from the descending gradient cluster, we calculated the mean T1w/T2w ratio across HCP subjects and the mean expression level of genes belonging to the descending gradient cluster in each cROI. We find that a given cytoarchitectonic region’s T1w/T2w ratio is positively correlated with the expression level of descending-gradient genes (r(13)=0.88, p=0.0001; Fig 4C, top).

Comparatively, ascending-gradient gene expression levels are positively correlated with cortical thickness across cROIs (r(13)=0.83, p<0.0001; Fig 4C, bottom). Cortical myelination is a factor that can influence *in vivo* estimates of cortical thickness from T1-weighted images, as more myelin in deeper layers of cortex near the gray-white boundary “whiten” those voxels at the boundary and shift the estimated boundary of cortex further towards the pial surface. To ensure that cortical thickness as measured from the HCP dataset is significantly correlated with ascending-gradient gene expression independent of myelination, we produced a stepwise regression model including both T1w/T2w values and ascending-gradient expression levels as predictors of cortical thickness. When both terms are included, ascending-gradient gene expression still accounts for a significant portion of thickness variation across cROIs (coefficient = 0.55, p=0.0004).

### Gene expression gradients emerge at unique points in human development

As genes are dynamically expressed not only across tissue compartments, but also throughout the lifespan, we asked at what point these gene gradients emerge during human development. To pinpoint the developmental trajectories of these gene expression gradients, we leveraged the developmental transcriptome of the BrainSpan^48^ atlas of the developing human brain (http://brainspan.org), which characterizes the expression magnitude of the same gene targets as in our previous analyses. While the cortical locations sampled in this dataset were fewer, early visual (peri-calcarine, which includes mostly hOc1) and late visual cortex (ventral temporal cortex or VTC, which includes the four FG areas) were sampled across 42 brains ranging from prenatal to adulthood. Though this coverage is sparser than that of the previous analyses, which covered all 13 areas of the visual hierarchy, it is (a) sufficient enough to compare the earliest (e.g. hOc1) vs. the latest (e.g. the four FG areas) stages of the hierarchy and (b) sensitive enough to detect a difference between these two stages as hOc1 and the four FG areas are the most differentiable based on their mean gene expression magnitudes (Fig 4). We hypothesized that opposing gene gradients (e.g. higher expression level in hOc1 compared to VTC and vice versa) would emerge prior to birth and then get stronger postnatally. To test this hypothesis, we first aimed to replicate our prior analyses by calculating the mean expression levels of the top 200 genes identified from the AHBA data in the 20-60 year old tissue samples from BrainSpan, which were closest to the age range used in the previous analyses in Figures 1-4. Illustrated on the far right of Figure 5, the ascending-gradient genes are expressed more in VTC compared to hOc1 (forming a positive slope) while descending-gradient genes are expressed most in hOc1 and least in VTC (forming a negative slope). As this replicated our main finding from the analyses in Figures 1-4, we further examined the expression patterns for ascending- and descending-gradient genes at earlier developmental timepoints. We performed an ANOVA on the expression slopes (the difference between VTC and hOc1 expression magnitudes) with grouping variables of gene cluster and developmental timepoint, revealing a significant interaction (F(1,34) =12.08,p<0.0001). Surprisingly, we find that those genes constituting the ascending gradient demonstrate a positive slope at the earliest gestational timepoint (10-12 post-conception weeks), but that those genes belonging to the descending gradient have yet to reach their adult-like expression pattern, and instead are expressed more highly in VTC than hOc1. It is not until 19-24 post-conception weeks that the descending-gradient slope diverges in sign from the ascending-gradient expression pattern. Furthermore, it is not until early childhood (before 5 years of age) that the two expression gradients achieve their adult-like mean expression values and characteristic crossover pattern.

**Figure 5:**
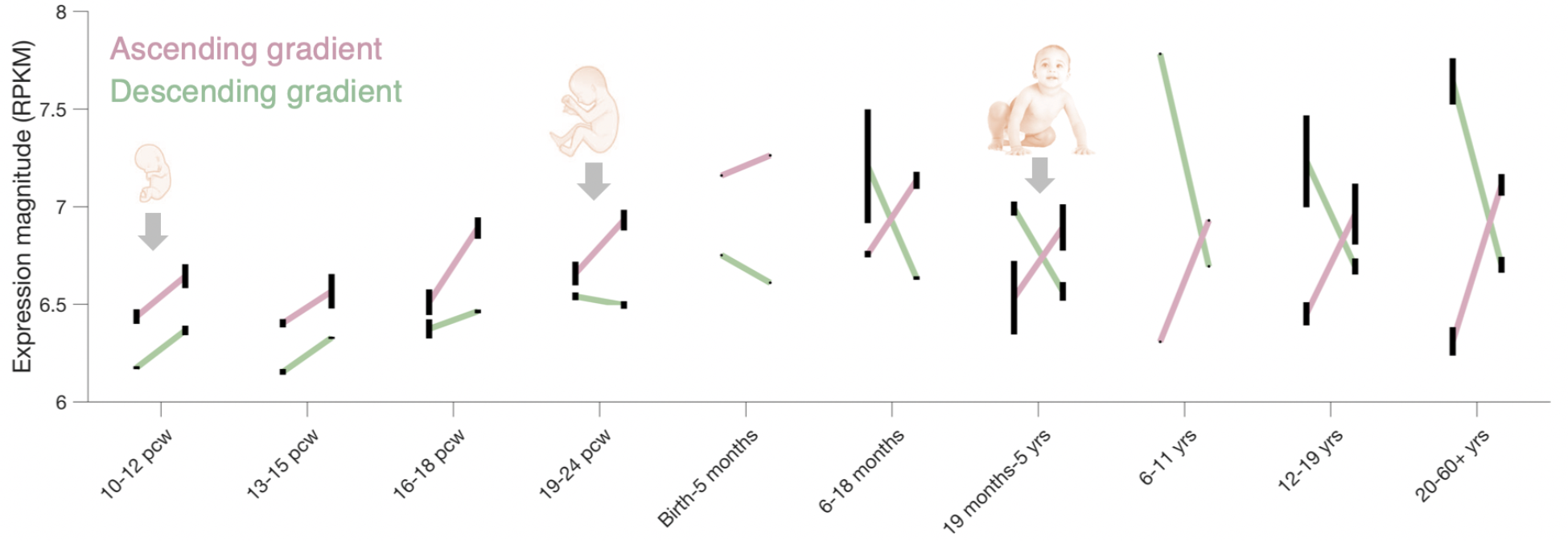
Ascending and descending gene gradients emerge at different points in human development. The two gene gradients derived from the top 200 differentially expressed genes in adults (ascending gradient clusters shown in pink and descending gradient clusters in green), show differences in their expression levels across development in early (left points in each line) and late (right points in each line) visual processing stages. At the earliest timepoint (10 post-conception weeks, pcw), the ascending gradient (pink) shows an increasing expression profile from early to late visual processing stages. However, the gradient genes that will show a descending expression profile (green) in adulthood show an increasing profile at this stage of development. It is not until 24 post-conception weeks that the two gene gradients diverge in slope. Errorbars reflect standard error across donor samples.

### Gene gradients emerge at unique points in evolution

The emergence of different cortical structures during gestation is thought to mirror their different evolutionary timepoints^49^. Indeed, a key distinction of humans from other primate species is the expansion of the cortical sheet^50,51^, whereby many structures extant in human high-level visual cortex, such as the fusiform gyrus and its constituent cytoarchitectonic regions, do not exist in the brains of non-human primates (NHP). These structures emerge later in the womb than those that are evolutionarily preserved with non-human primates - for example, the calcarine and parieto-occipital sulci^52^. To see if these genetic gradients contribute to the unique anatomical features of human versus non-human cortex, and to help pinpoint the evolutionary origin of the human visual processing hierarchy, we examined the expression pattern of the same ascending- and descending-gradient genes in the macaque monkey^53^ using a transcriptome analysis performed similarly to the AHBA and BrainSpan datasets. Like the BrainSpan data, the NHP dataset did not sample the cortex with the same density as the AHBA, but did sample early visual cortex (V1 and V2) in addition to an area in high-level visual cortex (area TE^54^), which is considered to be homologous to human VTC, and to which we refer to as “late” visual cortex for the present analyses (Materials and Methods). Taking a similar approach as the BrainSpan dataset and analyses (Fig 5), we hypothesized that should the two opposed gene clusters be conserved in the adult macaque, then tissue samples from late visual cortex should show higher expression magnitudes of ascending-gradient genes and lower expression magnitudes for descending-gradient genes, compared to early visual cortex. We find that while the genes identified in humans that form the ascending-gradient also form an ascending gradient in adult macaques, showing higher expression magnitudes in late compared to early visual cortex (Fig 6, right), the descending-gradient expression profile is not conserved, showing similarly high expression levels in late compared to early visual cortex (Fig 6, left). For both the ascending-gradient and descending-gradient genes, expression magnitudes are significantly higher (ascending: t(50)=4.53, p<0.0001; descending: t(50)=4.35, p<0.0001) in late compared to early visual cortex. Thus, our analyses show that the genes contributing to the ascending gradient of the visual processing hierarchy is evolutionarily preserved in macaques. Additionally, our analyses also show that humans have a series of genes contributing to the descending gradient of the visual processing hierarchy that is either a) not preserved in macaques, or b) also present in macaques, but guided by a different set of genes compared to humans, which we consider further in the discussion.

**Figure 6:**
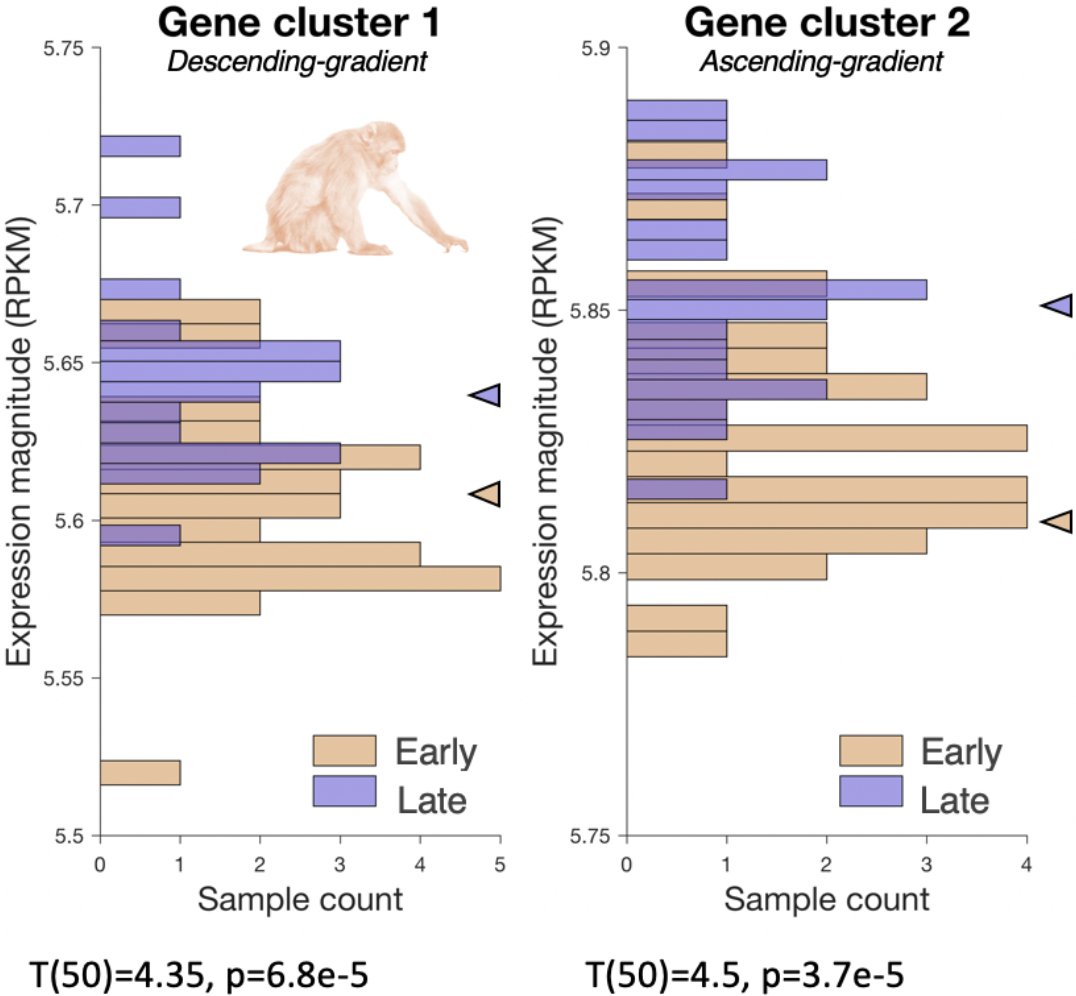
Ascending, but not descending, gradients contribute to the differentiation between early and late visual processing stages in macaques. Histograms of expression magnitudes from all tissue samples in early visual cortex (V1/V2, shown in tan) and late visual cortex (area TE, shown in purple). Histograms are shown separately for those genes that show a descending gradient in humans (gene cluster 1) and those genes that show an ascending gradient in humans (gene cluster 2). The average expression magnitude in the early and late cortical ROIs are denoted with triangles. RPKM = reads per kilobase million. The gene expression magnitude is higher in the late compared to the early processing stage for both gene clusters, which suggests that the descending gradient is not conserved across species and may underlie differences between the visual processing hierarchy across species.

## Discussion

Leveraging recent advances and shared datasets of human cytoarchitectonic areas in occipitotemporal cortex^7–13^ and maps of gene expression across cortex^42,48,53^, we demonstrate that the human visual system achieves its consistent anatomical layout and areal differentiation across individuals in cortex via two opposed, gene expression gradients. We find that the transcription magnitude of a sparse subset of human genes are capable of capturing not only the known hierarchy of human visual cortex, but also that the genes cluster into two classes: one whose expression magnitude increases as one ascends the visual hierarchy and correlates with the thickness of cortex, and another whose expression decreases along the hierarchy and correlates with the ratio of T1w/T2w values, which are related to cortical myelin content. When examining the developmental trajectory of these transcription gradients, we find that the ascending gradient emerges earlier in the gestational process during the first trimester, compared to the descending gradient which does not emerge until the second. Mirroring its delayed developmental emergence, the descending transcription gradient appears to be recent in primate evolution, as it is not conserved in the macaque visual system in which the homologous genes only form ascending gradients. Together, and to our knowledge, these findings not only establish one of the first multiscale models linking genetic expression, cytoarchitecture, and cortical folding in the human brain, but also help pinpoint the genetic underpinning of the cortical organizational principles underlying visual processing hierarchies in human cortex.

Furthermore, we would like to emphasize the importance of, as well as value in, being second. That is, our present findings replicate and extend the recent findings of Burt and colleagues^40^. Given the present empirical climate in which we as scientists are all embedded - one that often stresses a crisis regarding replication - we are pleased that our findings so quickly replicate those from Burt and colleagues. Importantly, we extend these findings, linking transcription to cytoarchitecture, human development, and the evolution of our species. In the sections below, we discuss our findings in the context of (a) a model linking genes, cells, and function in the human brain, (b) gradients as a principle of hierarchical cortical circuits, (c) development as a window into the origins of the brain, and (d) how the descending gradient may also be related to myelination and other macromolecular processes in hominoid-specific structures of the visual processing hierarchy.

### A model linking genes, cells, and function in the human brain

Taking advantage of updated definitions of cytoarchitectonic areas in human visual cortex, we demonstrate that areas differentiated by their cellular make-up are further distinguished by the extent to which they express two classes of genes. The advantage of using cytoarchitecture to define the hierarchical stages of visual cortex - rather than correlated proxies of hierarchical stages^40^ - is twofold. First, cytoarchitectonic divisions give us access to the underlying cellular organization that compose different processing stages of a cortical hierarchy, which ultimately enables us to link the expression levels of different genes to the organization of neural circuitry in greater detail. For example, one gene belonging to the ascending-gradient cluster is NECAB2, a protein-encoding gene involved in the intracellular scaffolding that responds to the presence of calcium, endowing cells with activity-dependent structural changes. The expression magnitude of NECAB2 is strongly under the influence of the gene PAX6^55^, a gene that is responsible for the proper structural organization of the eye, retina, and the inherited retinotopic organization of the cortex^56^. The multimodal model resulting from our findings provides a basis for understanding the origins of the visual system’s organization, suggesting that the retinotopic organization observed in even the highest-levels of the human visual system^34,57–59^ may be an inherited feature from a gene responsible for organizing the peripheral nervous system. While this may seem far-fetched, empirical findings provide strong support for this idea. For example, enucleation influences both the folding of the calcarine sulcus, as well as the cytoarchitectonic structure within it^60,61^.

The second strength of using cytoarchitectonic definitions of the visual system arises from work by our group demonstrating that cytoarchitecture is coupled with the folding of the cortex and with different functional regions. This allows our multimodal model linking genes, cytoarchitecture, and brain function to be extended into the brains of living subjects in which cytoarchitecture cannot be measured directly. This provides a powerful tool for future research to relate genetic mutations with specific brain regions. Surface-based tools allowing the alignment of individual brains with respect to the folding of cortex preserves structure-function relationships^25,62^, and will allow researchers to ask, with greater detail, how a given brain region maps both into 1) cytoarchitecture and 2) the genetic expression gradients described here. For example, the GABRB gene identified as belonging to the ascending-gradient cluster is highly expressed in the highest-levels of the visual processing hierarchy, including regions within the lateral Fusiform Gyrus (FG2, FG4) that contain face-selective regions, which are critical for face perception^30,43^. The GABRB gene and its related subtypes are protein-encoding genes involved in inhibitory GABA receptors located on post-synaptic dendrites, and have been consistently implicated in autism spectrum disorder^63^. Furthermore, given GABA receptors’ prominent role in cortical plasticity and establishing the structural connectivity of the cortex^64,65^, functional and structural abnormalities observed in the cortex of individuals with autism^66–69^ can now be linked with greater specificity in living subjects. Of course, we have only selected a subset of genes to illustrate how the results of the present study could influence future studies. Future work examining the roles of any of the 200 genes (Supplementary Table 1), especially those in the descending gradient, will greatly advance our understanding of the role that specific genes play in the functional or dysfunctional development of the human visual processing hierarchy.

### Gradients as a general principle of hierarchical cortical circuits

In addition to the observation that the visual processing hierarchy is organized along two genetically opposed expression gradients, we found that the expression magnitude of each gene cluster was tightly correlated with either the local cortical content of myelin (e.g. the T1w/T2w ratio) or the thickness of cortex. Through enrichment analyses^70,71^, we find that as a whole, the ascending-gradient gene cluster contains genes that primarily play a role in synaptic regulation and ion transport, implicating these genes in cell-to-cell signaling which likely underlies their tight relationship to the thickness of cortical layers. Indeed, previous research has linked the transcription of PPP4R4 (protein-encoding pyramidal cell marker) directly to cortical thickness in humans^72^. Descending-gradient genes, in contrast, play a strong role in macromolecule metabolic processes. Myelin is one of the most ubiquitous macromolecules in the human brain, and its decreased presence in the cortex as one traverses further from V1 into ventral temporal cortex is likely under the control of multiple genes within the descending-gradient cluster, given their strong correlation. Rather than attempt to link specific genes with either of these anatomical measures of the cortex, we instead emphasize that feature gradients appear to be a key hallmark of hierarchies in human cortex. In addition to likely driving the structural changes that result in the cytoarchitectonic differentiation of different stages of the hierarchy, genetic gradients undoubtedly play a major role in establishing functional gradients of the human visual system. For example, recent work has demonstrated that the ventral processing stream in humans is marked by a temporal processing gradient^73–75^ whereby increasingly high-level visual regions demonstrate longer temporal windows over which they integrate stimulus information. Furthermore, one of the first observations about neurons in the visual system was that their receptive fields—the portions of visual space in which a stimulus is capable of evoking a response—grow in size as one ascends the hierarchy of visual field maps^54,59,76^. Even *in silico* simulations of deep neural networks achieve high classification accuracy in object recognition tasks with neural architectures that employ gradients of increased pooling from one layer to the next^77,78^, suggesting that functional and structural gradients are likely a fundamental principle of processing hierarchies. Thus, genetic gradients as observed in the present study likely endow swaths of cortex with structural gradients (e.g. myelin, receptor densities, etc.) from which functional abilities emerge differentially across cortical regions that sample unique points along these gradients. Evidence for this possibility is further supported by the fact that the clustering of visual areas using cytoarchitectonic properties^10^ does not *exactly* match the clustering when using gene expression profiles. Thus, though two opposing gene expression gradients explain the clustering of the visual processing hierarchy (the arealization of which was defined using cytoarchitecture), the genes within both gradients likely also contribute to the endowment of these swaths of cortex with structural gradients from which functional abilities of neural circuits emerge.

One implication from the genetic gradients measured in the current study is that even a subtle single-nucleotide polymorphism in a single gene can have a widespread impact at every point along a cortical hierarchy. Indeed, autism spectrum disorder, despite being characterized often as a social disorder, has been documented to impact even early visual cortex and low-level visual behaviors^79,80^. Conversely, seemingly unrelated mutations in genes belonging to separate gradients could cause focal structural and functional impairments in a single brain region. For example, a cortical region that requires long integration times and unmyelinated local connectivity features that the current findings suggest are controlled by genes belonging to separate gradients may be uniquely impacted by single gene polymorphisms in each gradient. This cortical region, which would require a) high expression magnitude of an ascending-gradient gene and b) low expression of a descending-gradient gene, would be disproportionately impacted by these mutations compared to mid-level visual regions that do not require such polarized expression levels. Such a scenario may underlie specific developmental deficits like dyslexia^81^ or prosopagnosia^82,83^ in which individuals can often times present with a selective perceptual deficit in reading or face recognition abilities, respectively.

Notably, the original investigation into the transcriptional landscape of the prenatal brain^48^ revealed that some genes (e.g., FGF9) demonstrate a descending gradient while others (e.g., CPNE6) show an ascending gradient from temporal to frontal cortex. As these genes also formed gradients along our visual hierarchy, this suggests that certain genes may play more universal roles in cortical organization outside of visual cortex. Indeed, pinpointing those genes that form gradients generally across cortex^40,48,72^ and those that form gradients specific to different sensory modalities will help pinpoint the genetic correlates of functional differentiation in different processing streams.

### Development as a window into the origins of the brain

In adults, the shared and consistent organization of the visual processing hierarchy is likely based upon the shared existence of the two genetically opposed expression gradients that we have identified in the present study. Nevertheless, examining the developmental and evolutionary origins of these genetic gradients reveals differential emergences during human gestation and across species. Specifically, we found that the ascending-gradient genes are qualitatively present at ten weeks of gestation (and perhaps earlier), while the descending-gradient genes do not achieve their characteristic, decreasing expression values across the hierarchy until the middle of the second trimester. Both of the gene clusters do not reach quantitative maturity until middle or late childhood, as they did not demonstrate their adult-like crossover pattern of mean expression levels until the age range of 6-11 years old. This pattern of results holds two powerful implications. Firstly, the structural differences that make up the unique cytoarchitecture in each stage of the visual processing hierarchy is likely controlled primarily by the ascending-gradient genes, as these genes are already differentially expressed along the hierarchy within the first trimester. It may thus be that the organization of the visual processing hierarchy is to a large extent under the control of a sparse set of genes, specifically those that increase in magnitude from V1 to ventral temporal cortex. The later emergence of the descending-gradient genes may aid in further refining functional differences between cytoarchitectonically unique regions, especially within those regions located in macroanatomical structures that are hominoid-specific (which we expand further on in the next section). Secondly, both gene gradients are actively changing in expression magnitude until children are well into their school-age years, suggesting that while genes likely establish areal differences in this cortical hierarchy before birth, they may also be responsive to visual experience well after, with individual differences in viewing behavior potentially up or downregulating the expression of certain genes. Future research may examine which gradient genes are sensitive to experience, and explore the possibility that an inability to undergo experience-dependent changes in expression levels could be another potential source of developmental cortical deficits.

It is a commonly held belief that the point during gestation at which a cortical structure emerges reflects its evolutionary age, with more ancient structures emerging early in fetal development such as primary sensory regions and primary cortical folds preceding evolutionarily novel structures appearing later, such as tertiary cortical structures^49,84,85^. Consistent with this hypothesis, we observed that the descending-gradient genes, which lag behind ascending-gradient genes during gestation, were not conserved in their expression pattern within the macaque visual cortex. These findings have several important implications. First, while the macaque is often used as a homologous species for studying the visual system,^40,86–90^ the known differences in the localization, presence, and organization^91–97^ of the visual systems across species may be partly explained by the same genes forming a descending gradient in humans, but not in macaques. Future research can explore yet-discovered functional differences between the visual systems of the two species, specifically as they pertain to structures encoded by those genes belonging to the descending gradient (see Supplementary Table 1). Another important implication of these findings is that they help pinpoint when certain cortical structures emerge in the evolutionary timeline. For example, as we demonstrate that genetically opposed gradients likely contribute to the visual processing hierarchy, the existence of additional cortical structures in humans—the Fusiform Gyrus^98^, a separate fourth ventral field map^28,99^, and larger cortical volume dedicated to mid- and high-level visual regions relative to V1^2,93,94,100^—may have emerged as a result of additional genetic gradient information. Additionally, it may be the case that the unique opposed-gradient pattern in humans is a factor that may contribute to unique human faculties. Finally, it may be the case that macaques and other species also have a descending gradient, but this gradient develops under the guidance of different genes compared to humans. This intriguing possibility can be addressed in future work, which, if found to be true, would shed light on what is similar and different regarding the roles that similar and different genes play in the development of the visual processing hierarchy across species.

### The descending gradient may also be related to myelination and other macromolecular processes in hominoid-specific structures of the visual processing hierarchy

An interesting conundrum produced from the present findings is that the descending gradient, which could not be identified in macaques, is highly correlated with myelination, as well as linked to other macromolecular processes based on enrichment analyses. And yet - which is perhaps likely obvious to the reader - macaque brains also have myelin and areas in the macaque brain have been delineated based on myeloarchitecture^14,101,102^. We propose that the descending gradient may be particularly critical for myelination (and other macromolecular processes) in cortical regions located in macroanatomical locations not present in macaques. Consistent with this proposal is the fact that the descending gradient does not emerge until 19-24 weeks of gestation, which is coincidentally a similar timepoint during which the human brain begins forming macroanatomical structures such as the Fusiform Gyrus (FG), which is only present in humans and non-human hominoids, but not macaques. Indeed, the FG emerges between 24-27 weeks, while tertiary sulci in the occipitotemporal lobe do not emerge until much later, at 40-44 weeks^84^. Thus, given the delayed emergence of the descending gradient in humans and its complete absence in macaques, it is likely that a subset of genes in the descending gradient play a role in myelination and other macromolecular processes associated with the arealization of regions within the visual processing hierarchy that are located within hominoid-specific macroanatomical structures.

### Conclusion

Combining cytoarchitectonic definitions of a human cortical processing hierarchy with genetic transcription analyses of the cortex, we produced one of the first (to our knowledge) multimodal models linking structure and function across a wide range of spatial scales in human visual cortex. Not only do these findings help establish the genetic basis of areal differentiation and the consistency of brain organization across individuals, but they also identify a sparse subset of genes that form opposing expression gradients across the human visual processing hierarchy. Furthermore, these cortical transcription gradients undergo differential development during human gestation, whereby genes form an ascending gradient earlier than a descending gradient. Lastly, we find that the opposed gradient pattern formed by these genes in humans is not conserved across species, with the same genes only showing ascending gradients in macaque visual cortex. These findings provide essential groundwork for understanding both the origins of cortical processing networks and the factors that can contribute to their maldevelopment, as our surface-based approach linking multiple measurement modalities within the human cortex enables future studies to link genetic analyses *in vivo* with transcriptomic, cytoarchitectonic, structural, and functional measurements all within the same individual.

## Materials and Methods

### Data

All data analyzed in the present manuscript were curated from 5 freely available datasets that were acquired, shared, and approved according to the Ethics Committees of each institution:

1. JuBrain: http://www.fz-juelich.de/JuBrain/EN/_node.html;
2. AHBA: http://brain-map.org/;
3. Human Connectome Project: http://www.humanconnectome.org/study/hcp-young-adult/data-releases
4. BrainSpan atlas: www.brainspan.org;
5. NIH Blueprint Non-human Primate (NHP) Atlas website: http://www.blueprintnhpatlas.org/;

#### 1. JuBrain cytoarchitectonic atlas and ROI definition

The JuBrain cytoarchitectonic atlas consists of a set of cytoarchitectonic brain areas that are defined by an observer-independent analysis of laminar cell density profiles in ten human postmortem brains (http://www.fz-juelich.de/JuBrain/EN/_node.html). Each area is represented by a 3D map that describes the maximum probability with which a certain cytoarchitectonic area can be assigned to a certain macroanatomical location in the brain (Eickhoff et al., 2005; Zilles and Amunts 2010). Presently, 100 areas have been mapped in the atlas, which covers about 70% of the brain. Among them, 13 visual areas located in occipitotemporal cortex have been delineated including hOc1, hOc2, hOc3d, hOc3v, hOc4d, hOc4v, hOc41p, hOc41a, hOc5, FG1, FG2, FG3, and FG4 (Figure 1). These 13 cytoarchitectonic regions of interest (cROIs) are the focus of the present study. As the JuBrain cytoarchitectonic atlas has been aligned to the MNI305 (Colin 27) space and the Allen Human Brain Atlas (AHBA) has been aligned to the MNI152 space, the JuBrain cytoarchitectonic atlas was first linearly transformed into the MNI152 space using FMRIB's Linear Image Registration Tool (FLIRT) to align the cROIs to the AHBA.

#### 2. Human transcriptome data and analysis

##### Allen Human Brain Atlas

The gene expression data used in the present study were obtained from the Allen Human Brain Atlas (AHBA), which is a publicly available atlas of gene expression and anatomy. The AHBA employs DNA microarray analyses to map gene expression from tissue samples taken broadly across human cortex. The dataset is based upon the conglomeration of measurements from 6 postmortem human brains, although not all brains were sampled identically. In the end, cortical samples were acquired using macrodissection from every brain and submitted to normalization analyses to make measurements within and between brains comparable. Each sample was associated with a 3D coordinate from its donor’s MRI volume and its corresponding coordinate (x,y,z) in the MNI-152 space. Each tissue sample was analyzed for the expression magnitude of 29,131 genes, with 93% of known genes being queried by at least 2 probes. For more details regarding how the microarray data were normalized, please see the following documentation: http://help.brain-map.org/display/humanbrain/documentation/.

##### Gene expression preprocessing

The collection and quality control of the gene expression data has been described previously (see online documentation link above). Here, we discuss the preprocessing of data for the study at hand. The raw microarray expression data for each of the six donor brains included the expression level of 29,131 genes profiled via 58,692 microarray probes. We implemented five preprocessing steps. First, probes were excluded that did not have either a) a gene symbol or b) an Entrez ID. This resulted in 20,737 genes. Second, the expression profiles of all the probes targeting the same gene were averaged. Third, in order to remove variability in gene expression across donors, gene expressions were normalized by calculating z-scores separately for each donor. Fourth, and as described above, samples were assigned to cROIs based on MNI152 coordinates. Specifically, samples were assigned to cROIs based on the Euclidean distance between the sample and a cROI. Fifth, because previous studies did not identify significant interhemispheric transcriptional differences, data from both hemispheres were combined. As a result, a total of 331 samples were mapped to the 13 cROIs in occipitotemporal cortex across all donors. The number of samples varied across cROIs (see Figure 1).

##### Gene selection

In order to identify the genes that were expressed differently across the visual hierarchy, we first divided the 13 cROIs into groups with equal samples. This was done in order to equalize (as close as possible) the number of samples in a given group for the purposes of statistical analysis. We divided the samples from the 13 cROIs into 4 groups: hOc1, [hOc2 and hOc3], [hOc4d, hOc4v, hOc41p, hOc41a], and [FG1, FG2, FG3 and FG4]. hOc5 was excluded because only 2 samples were included within that region of cortex. Once the cROIs were divided into groups, for each gene, we ran a 1-way ANOVA with cROI group as a factor. We ranked the genes according to the p-values from the ANOVA and selected the top ~1% (200) of genes for further analysis. This thresholding approach was taken in order to avoid including genes that may result in significance simply as a function of multiple comparison. The names of these 200 genes are listed in Table 1 and colored according to gradient membership.

To ensure that our results were not limited to the method used to select the genes, we repeated our analyses using a principal component analysis (PCA) on the full gene expression data from the 13 cROIs. As illustrated in Supplementary Figure 1, our results are not dependent on the method used to select the genes. With either method, two opposing gradients are identified. With the PCA approach, the two gradients represent positive and negative directions on the first PC (PC1 in Supplementary Figure 1), which explains 33.8% of the variance. The PCA is described in detail in Supplementary Figure 1.

**Supplementary Table 1:**
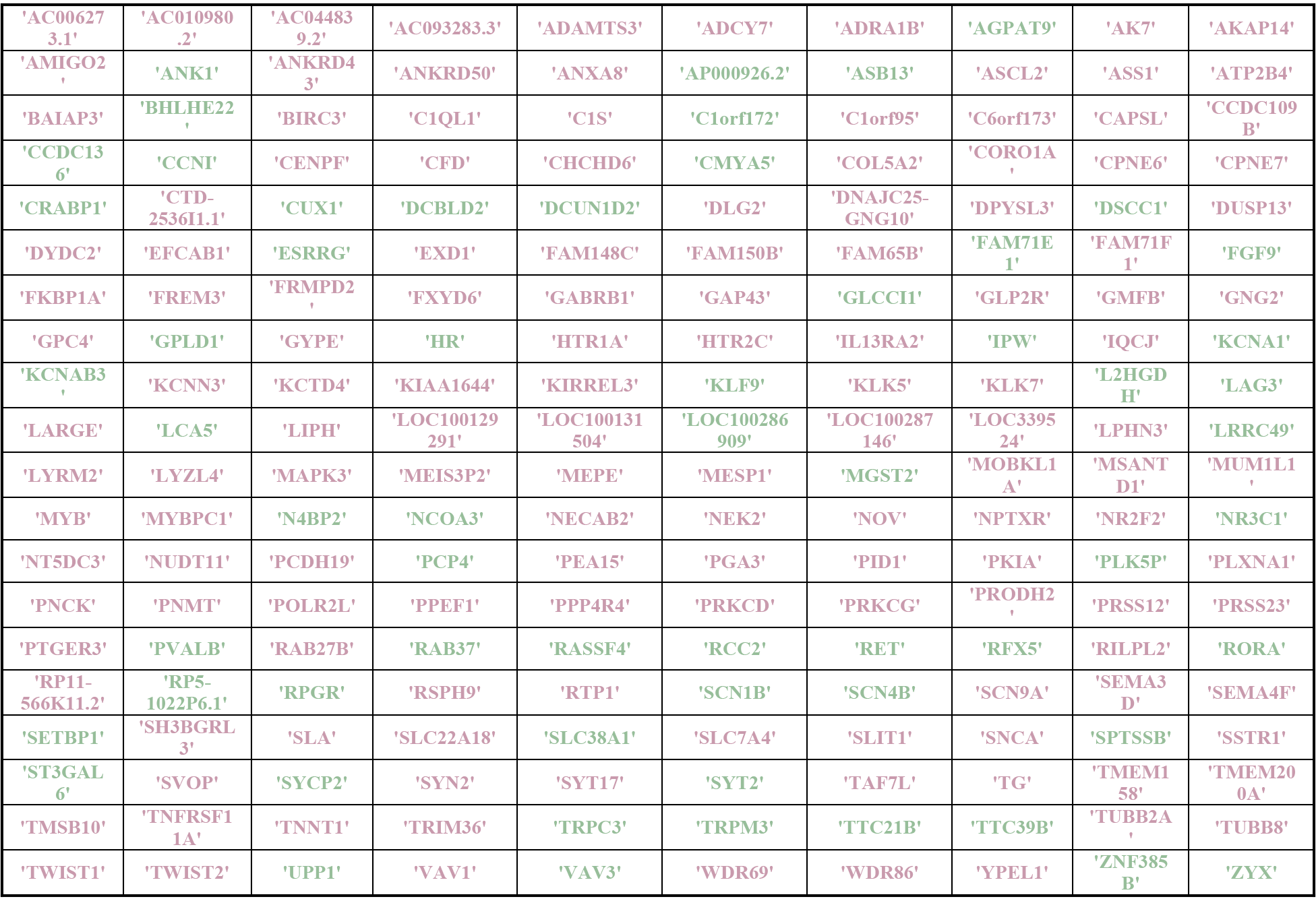
Gene symbols of the top 200 genes. Genes written in green belong to the descending gradient cluster, while those in pink belong to the ascending gradient.

##### Exploring the relationship of gene expressions among cROIs

In order to further explore the pattern of gene expression across the cROIs of the visual processing hierarchy, the microarray data were first averaged across all samples from all donors in the matched cROI across both hemispheres. Accordingly, a 200 × 13 data matrix was generated, which represents the expression pattern for the top 200 genes across the 13 cROIs. We then used this matrix as the input to an agglomerative hierarchal analysis using Euclidean distances and the weighted-pair group method with arithmetic mean (WPGMA) in order to examine the relationship of gene expression among the cROIs of the visual processing hierarchy. Prior to clustering, the 200×13 matrix was sorted according to each cROIs Euclidean distance from the expression pattern of hOc1. In this way, the ordering of cROIs along the x-axis in the resulting dendrogram is meaningful, as it represents at the level of dendrogram leaves the distance from hOc1. As a result, this dendrogram has rooted leaves and the clusters at this lowest level are not rotatable as in other dendrograms. To evaluate the significance of this dendrogram ordering, we evaluated how often such an ordering of these region can occur by chance. To accomplish this, we utilized a bootstrap approach with 10,000 iterations. On each bootstrap, within each cROI we randomly shuffled the expression magnitudes of the 200 genes before submitting the cROI expression profiles to the agglomerating clustering algorithm. We then took the resulting ordering vector from the clustering algorithm (e.g., [2 11 5 3…7]) and calculated its Euclidean distance from the true ordering vector (e.g., [1 2 3 4 5…13]). We obtained the p-value as the resulting probability of observing a zero Euclidean distance from the resulting 10,000 samples.

#### 3. Human Connectome Project measures of T1, T2, and cortical thickness

##### Data

Cortical maps of T1-weighted intensity, T2-weighted intensity, and cortical thickness were obtained from the Human Connectome Project^103^ which included 1096 participants. In each subject, T1- and T2-weighted scans were collected using 3-Tesla MRI. While not a direct measure of cortical myelination, the ratio of T1-weighted to T2-weighted scans removes MR-related image inhomogeneities to a give voxelwise signal that is sensitive to the influence of myelin and iron (which are strongly co-localized in cortex). Measures of cortical thickness were produced using the automatic tissue segmentation algorithm in FreeSurfer^46^.

##### ROIs

As these measurements were projected to individual HCP cortical surfaces, we transformed cROIs into each individual HCP subject using cortex-based alignment (CBA) by first transforming the cROIs to the FreeSurfer average surface and then transforming the cROIs into individual HCP surfaces to ensure that individual differences in cortical folding were respected in each subject. This methodology assures that the coupling between cytoarchitecture and cortical folding observed in the postmortem brains is maintained in living subjects^25,30,43^.

##### Analyses

After aligning cROIs to each individual HCP participant, we then extracted the mean T1w/T2w value and mean cortical thickness value within each cROI for all 1,096 HCP subjects Distributions of these means across subjects for all cROIs is illustrated in the violin plots of Figure 4A. We statistically examined the relationship between anatomical gradient (T1w/T2w, thickness) and cROI in two ways: (1) with an analysis of variance (ANOVA; Fig 4A) and (2) the correlation between anatomical gradient and mean gene expression value (Fig 4C).

#### 4. Developmental transcriptome data and analysis

##### Data

The BrainSpan atlas contains gene expression data taken from cortical structures spanning 13 stages of human development across 8-16 brain structures. The atlas queried the expression of 17,604 genes via DNA microarray analyses similar to AHBA described above. We focused our analyses on transcriptome data from two ROIs: primary visual cortex and ventral temporal cortex (VTC). Primary visual cortex for intrauterine cases included samples taken in the vicinity of what would become pericalcarine cortex, including mostly V1. VTC was taken from the region of cortex that would become inferior temporal cortex which corresponds roughly to the four cytoarchitectonic ROIs labeled FG1-4 in adult cortex (Fig 1). As these areas cannot be recognized before 10 post-conceptional weeks (pcw), the developmental data from the first two timepoints (1 and 2A) were excluded from further analyses. The gradient genes we identified from the AHBA dataset (e.g., top 200) were then targeted for our analyses. See Supplementary Table 2 for the stage division and the corresponding number of samples. Information about data preprocessing and normalization is available from the BrainSpan Atlas website (http://help.brain-map.org//display/devhumanbrain/Documentation).

**Supplementary Table 2.** The definitions of developmental stages and the corresponding number of samples in each stage.

| Stage | Age | Developmental Period | Number of samples (V1, VTC) |
| --- | --- | --- | --- |
| 1 | 4-7 pcw | Embryonic | Not used |
| 2A | 8-9 pcw | Early prenatal | Not used |
| 2B Early | 10-12 pcw | prenatal | 3,3 |
| 3A | 13-15 pcw | Early mid-prenatal | 2,3 |
| 3B | 16-18 pcw | Early mid-prenatal | 4,4 |
| 4 | 19-24 pcw | Late mid-prenatal | 4,4 |
| 5 | 25-38 pcw | Late prenatal | 1,2 |
| 6 | Birth-5 months | Early infancy | 1,1 |
| 7 | 6-18 months | Late infancy | 2,2 |
| 8 | 19 months-5 yrs | Early childhood | 4,4 |
| 9 | 6-11 yrs | Late childhood | 1,1 |
| 10 | 12-19 yrs | Adolescence | 3,3 |
| 11 | 20-60+ yrs | Adulthood | 6,6 |

##### Analyses

Using these data, we first validated if the observed spatial gradients (ascending and descending) that we identified in the AHBA dataset could be replicated from the adult timepoint in the BrainSpan dataset. Once we verified that the ascending and descending gradient genes demonstrated their expected expression levels in V1 and VTC, we then measured their expression magnitudes in the same ROIs in early timepoints (Fig 5). At each timepoint, we averaged gene expression (reads per kilobase million) across all samples and donors within a given ROI to obtain an average ascending or descending gradient expression magnitude by timepoint.

#### 5. Non-human primate (NHP) transcriptome data and analysis

##### Data

The DNA microarray data for the non-human primate (NHP) were from the “Transcriptional Architecture of the Primate Neocortex” study^53^, a sub-project of the NHP atlas. Samples were collected from 10 cortical areas using laser microdissection in four adult rhesus monkeys. Each area was assayed using Affymetrix GeneChip Rhesus Macaque Genome Arrays (19, 050 genes; 52, 865probes). For the present analyses, transcriptome data from three ROIs were considered: V1, V2, and TE (which is considered a homologue of human VTC). For the sake of our analyses, V1 and V2 were grouped into a single ROI, which we refer to as *early visual cortex* in Figure 6. We also refer to data from TE as *late visual cortex* in Figure 6. The expression profiles of all the probes representing one gene were averaged within an ROI and across monkeys. Sample numbers can be seen in the histograms of Figure 6. As the cortex was sectioned via laser microdissection, sections from individual cortical layers were submitted to microarray analysis separately. In order to maximize the homology of analyses between humans and macaques, we averaged the data across all six layers in macaques. More complete descriptions of the experimental and data processing methods are available in protocol documents at the NIH Blueprint NHP Atlas website: (http://www.blueprintnhpatlas.org/)

##### Analyses

We first took the top 200 genes identified in adult humans and identified their NHP homologue using a gene-symbol-mapping table provided by the NHP atlas. 153 human genes had a match in the macaque gene set; the other 47 were likely not measured. 36 of these missing genes belonged to the ascending gradient (the larger of the two gradients), and 11 belonged to the descending gradient. The expression magnitude (RPKM) of all the probes representing a single gene were averaged together. Rather than summarize each ROI’s expression with a single mean across tissue samples, we instead created histograms illustrating the average expression magnitude of either descending or ascending gradient genes (Fig 6) to better characterize the distribution across tissue samples.

---

**Supplementary Figure 1:**
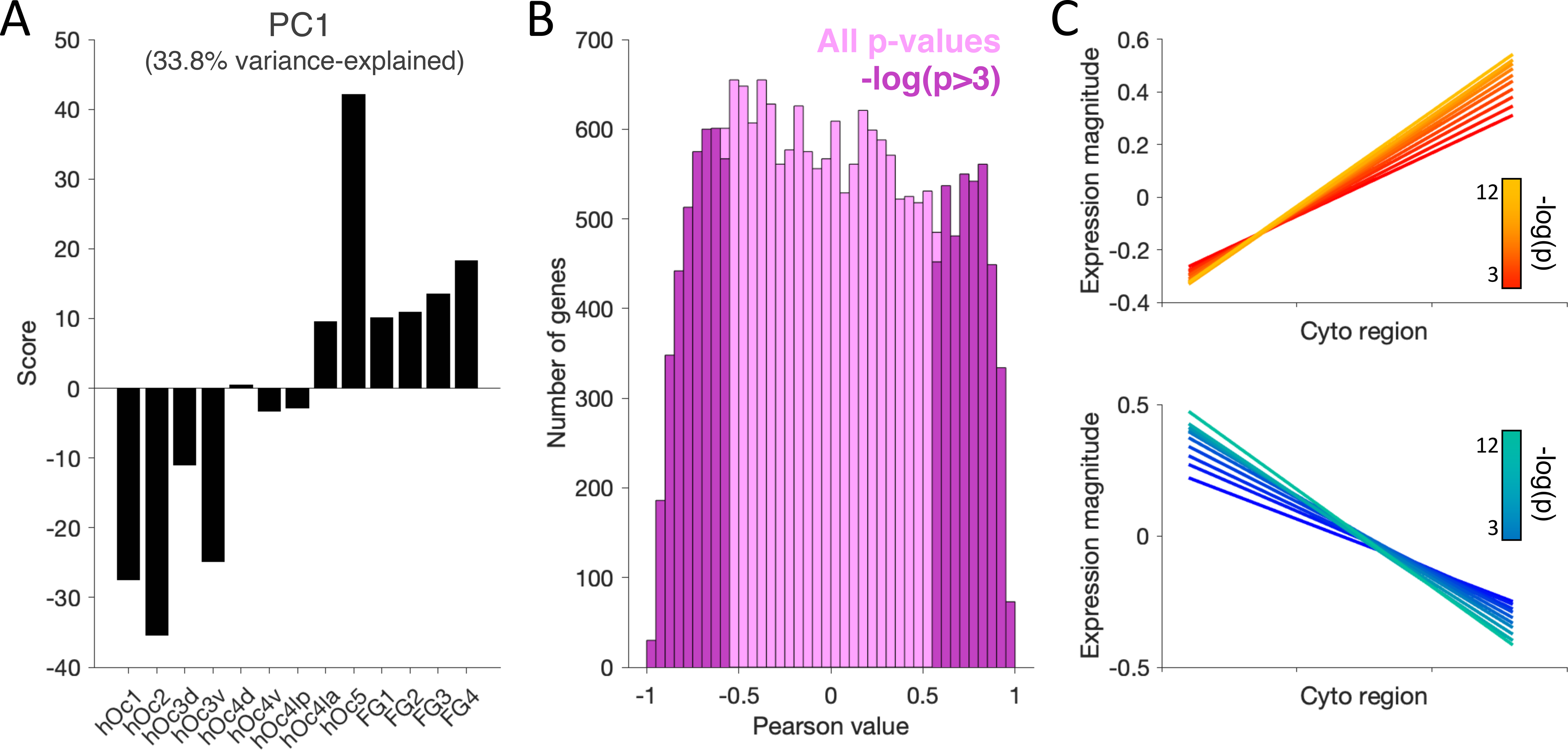
Principal component analysis (PCA) of averaged transcription profiles captures the ascending and descending gradient pattern in the first component. (A) We submitted the average expression magnitude of all genes within each of the 13 cROIs to PCA, the first component of which demonstrates a gradient of weighting scores, with early visual cortex mapping negatively onto this PC and later visual regions mapping positively. (B) Histogram of Pearson correlation significance values (negative log transformed) between each individual gene’s expression magnitude across the 13 cROIs with the scores of the first PC. Those genes with -log p-values exceeding 3 are highlighted in darker pink. (C) Linear fits summarizing the expression magnitude across cROIs of genes demonstrating either a positive (warm colors, top) or negative (cool colors, bottom) correlations with PC1 scores. We took a stepwise approach, first including all genes whose -log p-values exceeded 3 in the average, and then incrementally increasing the threshold until we only included genes in each group whose p-values exceeded 12, which included approximately 200 genes, equivalent to the group of 200 genes we chose in Figure 1. The magnitude of the positive or negative slope describing the expression gradients across cROIs increases as one includes more significantly differentially expressed genes. These analyses reveal that our results are not dependent on the method used to select the genes. With either method, two opposing gradients are identified. With the PCA approach, the two gradients represent positive and negative directions on the first PC, which explains 33.8% of the variance.

## Acknowledgements

This work was supported by (1) start-up funds provided by the University of California, Berkeley and the Helen Wills Neuroscience Institute (KSW), (2) Ruth L. Kirschstein National Research Service Award F31EY027201 (JG), and (3) the National Natural Science Foundation of China 31771251 (ZZ). We thank Jack Gallant, Kendrick Kay, Kalanit Grill-Spector, and Aviv Mezer for useful discussions regarding this work, and Evgeniya Kirilina for histological images.

